# Chloride intracellular channel (CLIC) protein function in S1P-induced Rac1 activation requires membrane localization of the C-terminus, but not thiol-transferase nor ion channel activities

**DOI:** 10.1101/2025.01.22.634370

**Authors:** De Yu Mao, Jordan J. Jesse, Daniel D. Shaye, Jan Kitajewski

## Abstract

We have established a novel and evolutionarily-conserved function for chloride intracellular channel proteins (CLICs) in regulating Rho/Rac GTPases downstream of G protein-coupled receptors (GPCRs). Endothelial CLIC1 and CLIC4 are rapidly and transiently re-localized from the cytoplasm to the plasma membrane in response to the GPCR ligand sphingosine-1-phosphate (S1P), and both CLICs are required to activate Rac1 in response to S1P, but how they perform this function remains unknown. Biochemical studies suggest that CLICs act as non-specific ion channels and/or as glutathione-S-transferases, dependent on N-terminal features, *in vitro*. Here we investigate CLIC functional domains and membrane localization requirements for their function in S1P-mediated Rac1 signaling. Structure-function analyses of CLIC function in endothelial cells demonstrate that CLIC1 and CLIC4-specific functions reside at their C-termini, and that the CLIC4 N-terminus encodes determinants required for S1P-induced re-localization to the plasma membrane but is dispensable for S1P-induced Rac1 activation when the C-terminus is localized to the plasma membrane via a heterologous signal. Our results demonstrate that the postulated ion channel and thiol-transferase activities of CLICs are not required for Rac1 activation and suggests that sequences in the CLIC C-termini are critical for this function. Given the importance of S1P signaling in vascular biology and disease, our work establishes a platform to further our understanding of the membrane-localized proteins required to link GPCR activity to Rho/Rac regulation.

## Introduction

Chloride intracellular channels (CLIC) are a family of evolutionarily conserved proteins whose physiological and molecular functions remained poorly defined until recent work from us, and others, demonstrated a role for them in G protein-coupled receptor (GPCR) and Rho-family GTPase signaling that is conserved from *C. elegans* to humans (Tavasoli et al., 2016; Mao et al., 2021; Arena et al., 2022; Kleinjan et al., 2023). CLICs can exist as nuclear, cytoplasmic, and/or membrane-associated forms, and different extracellular signals or physiological conditions can promote re-localization between these different cellular compartments (Novarino et al., 2004; Ponsioen et al., 2009; Malik et al., 2010; Gurski et al., 2015; Lecat et al., 2015; Carlini et al., 2020; Mao et al., 2021; Kleinjan et al., 2023). Although some CLICs transiently, or constitutively, reside at the plasma membrane, chloride channel activity for these proteins under physiological conditions has been challenging to establish (reviewed in Argenzio and Moolenaar, 2016), raising the question as to whether their postulated ion channel activity is required for downstream functions. Moreover, the crystal structure of invertebrate and human CLICs (Littler et al., 2004; Littler et al., 2005; Cromer et al., 2007; Littler et al., 2008) revealed that these proteins resemble the omega family of glutathione-S-transferases (Ω-GSTs), and purified CLIC1 and CLIC4 exhibit GST activity *in vitro* (Al Khamici et al., 2015). However, to date it is not known if the proposed GST and/or ion channel activities are relevant to CLIC function in GPCR and Rho-family GTPase signaling.

Out of six mammalian paralogs, CLIC1 and CLIC4 are expressed in the endothelium (Tung et al., 2009; Tung and Kitajewski, 2010; Mao et al., 2021), and we previously used human umbilical vein endothelial cells (HUVEC) to establish that CLIC1 and CLIC4 are required for cell viability, migration, and angiogenic behaviors (Tung et al., 2009; Tung and Kitajewski, 2010). We also found that activation of the sphingosine-1-phosphate (S1P) and thrombin GPCR pathways induces transient re-localization of CLIC1 and/or CLIC4 to the plasma membrane, and that CLIC1 and CLIC4 function in distinct GPCR cascades to regulate RhoA and/or Rac1 activation (Mao et al., 2021; Kleinjan et al., 2023). Specifically CLIC1 and CLIC4 function in sphingosine-1-phosphate receptor (S1PR) signaling (Mao et al., 2021) *in vitro*, while CLIC4 functions in the thrombin/protease activated receptor (PAR) pathway not only in cultured endothelial cells, but also *in vivo* (Kleinjan et al., 2023). Here evaluate CLIC function in S1P signaling (summarized in Fig. 1A). S1P binding to S1PR1 activates its coupled Gα subunit, Gα_i_, which in turn activates Rac1, promoting cell spreading, migration, and endothelial barrier enhancement (Sugimoto et al., 2003; Kono et al., 2004; Reinhard et al., 2017), while S1P binding to S1PR2 and S1PR3 activates RhoA, via Gα_12/13_, leading to stress-fiber formation, cell contraction and endothelial barrier disruption (Lee et al., 1999; Miura et al., 2000; Blaho and Hla, 2011; Reinhard et al., 2017). We found that S1P induces a rapid and transient re-localization of both CLIC1 and CLIC4 from the cytoplasm to the plasma membrane, and that both CLICs are required for S1P-induced Rac1 activation, while only CLIC1 is required for RhoA activation (Mao et al., 2021). Notably, CLIC1 and CLIC4 have non-overlapping functions in S1P-induced RhoA and Rac1 activation, as CLIC1 overexpression does not rescue the Rac1 activation defect caused by RNAi-mediated *CLIC4* knockdown, nor does CLIC4 overexpression rescue the Rac1 or RhoA activation defects caused by *CLIC1* knockdown (Mao et al., 2021).

**Fig. 1:**
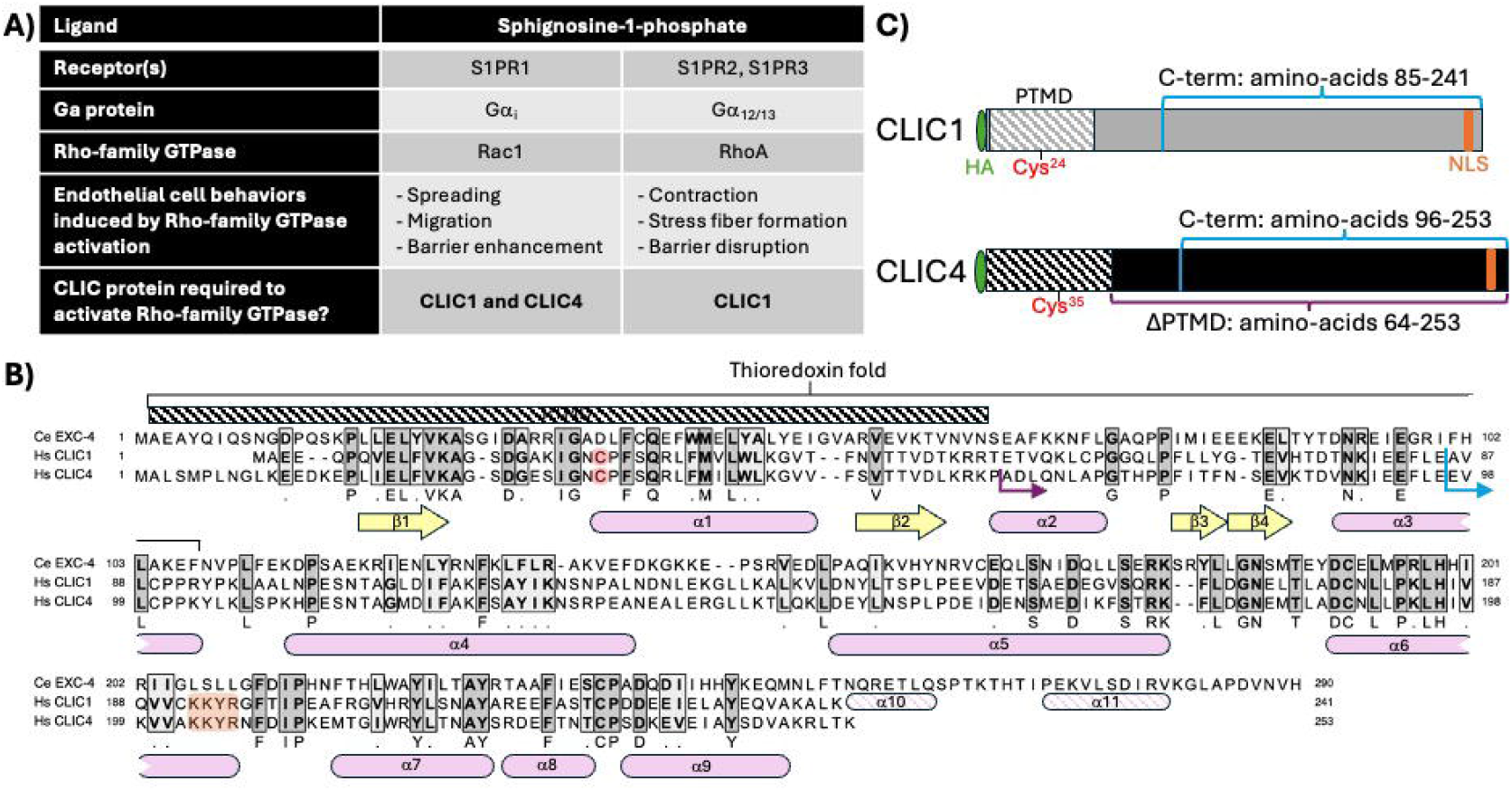
CLIC1 and CLIC4 in endothelial S1P signaling, and CLIC sequence and structural features relevant to this study. (**A**) description of shared and unique functions for CLIC1 and CLIC4 in endothelial S1P signaling (Mao et al., 2021). (**B**) Protein sequence alignment of *C. elegans* EXC-4, and human CLIC1 and CLIC4 indicating regions of amino-acid identity (dark grey shading) and similarity (light grey shading). Conserved beta sheets (yellow arrows) and alpha helices (pink rounded boxes) determined by crystal structure of these proteins (Littler et al., 2004; Littler et al., 2005; Littler et al., 2008) are shown. The N-terminal putative transmembrane domain (PTMD) and conserved thioredoxin fold are highlighted. Note that the N-terminal cysteine required for thiol-reductases activity (red), and the C-terminal nuclear localization signal (orange) found in human CLIC1 and 4 are not present in *C. elegans* EXC-4, while the *C. elegans* protein has a C-terminal extension with additional alpha helices (α10, α11). (**C**) cartoon representation of HA-tagged full-length CLIC1 and CLIC4 constructs. The C-terminal fragments swapped in the chimeric constructs analyzed in Fig. 2 are denoted in blue, and the ΔPTMD CLIC4 truncation analyzed in Figs. 3 and 4 is denoted in purple (the precise protein sequence location for these constructs are shown as blue and purple arrows in panel B).

Function of the *C. elegans* CLIC ortholog EXC-4 is strictly dependent on membrane localization (Berry et al., 2003; Berry and Hobert, 2006), and S1P induces rapid re-localization of CLIC1 and CLIC4 to the plasma membrane (Mao et al., 2021). The goals of this study are to determine which domain(s) mediate CLIC1 and CLIC4-specific functions in GPCR-induced Rho-family GTPase activation, and if these domains function at the plasma membrane. We demonstrate that the CLIC1 and CLIC4 C-termini define their specific functions in Rac1 activation in response to S1P in endothelial cells. Moreover, we show that the CLIC4 N-terminus is required for S1P-induced re-localization from the cytoplasm to the plasma membrane, consistent with the previously-described function for an N-terminal putative transmembrane domain (PTMD) in membrane accumulation of EXC-4 in *C. elegans* (Berry et al., 2003; Berry and Hobert, 2006). Finally, we show that the CLIC4 N-terminus is dispensable for S1P-induced Rac1 activation and endothelial barrier function when replaced with a heterologous plasma membrane-targeting motif; the myristylation (myr) signal of the kinase Lck (Bijlmakers et al., 1997; Zlatkine et al., 1997). Given that the myr signal does not possess GST activity or form ion channels, our results demonstrate that these postulated CLIC activities are dispensable for their function in GPCR-induced Rho-family GTPase activation. Our results support a model where CLIC C-termini function at the membrane to facilitate GPCR regulation of Rho-family GTPases.

## Results

### CLIC1 and CLIC4 functions in S1P-induced Rac1 activation are specified by their C-termini

Structure-function studies in *C. elegans* showed that the first 66 N-terminal amino-acids of EXC-4 are necessary and sufficient for plasma membrane localization (Berry et al., 2003; Berry and Hobert, 2006), defining a putative transmembrane domain (PTMD) that encompasses two ß-sheets flanking an α-helix; structural elements that are conserved in CLIC1 and CLIC4 (Fig. 1B. Harrop et al., 2001; Littler et al., 2005; Littler et al., 2008). The PTMD resides within the “thioredoxin fold” (Fig. 1B), which encodes the thiol-transferase activity of Ω-GSTs that is dependent on a catalytic cysteine found at the start of the first alpha helix (α1, red in Fig. 1B, C). The overlap between the PTMD and the thioredoxin fold, combined with biochemical and biophysical studies (Littler et al., 2004; Singh and Ashley, 2006; Goodchild et al., 2009; Goodchild et al., 2010; Al Khamici et al., 2015; Hare et al., 2016), has led to a model where redox conditions and GST-like activity of CLICs induces a conformational change that exposes the PTMD, resulting in plasma membrane insertion of the N-terminus of CLIC proteins. However, the cysteine required for GST activity is absent from *C. elegans* EXC-4 (Fig. 1B), which constitutively localizes to the plasma membrane (Berry et al., 2003) and which we showed also regulates Rho/Rac signaling in *C. elegans* (Arena et al., 2022). Therefore, questions remain as to whether the postulated GST or channel activities are indeed required for EXC-4/CLIC function in Rho-family signaling,

We previously established functional assays for CLIC1 and CLIC4 by infecting HUVEC with lentivirus expressing short-hairpin RNA to reduce *CLIC1* or *CLIC4* expression (hereafter *CLIC1*^*KD*^ or *CLIC4*^*KD*^) and co-infecting with lentivirus expressing N-terminally HA-tagged CLIC1 or CLIC4 (see Methods and Fig. 1C). Using this approach, we established that HA-CLIC1 rescued phenotypes caused by *CLIC1*^*KD*^ but not *CLIC4*^*KD*^, while HA-CLIC4 rescued *CLIC1*^*KD*^ but not *CLIC4*^*KD*^ (Mao et al., 2021), indicating that there is specificity to CLIC function in S1P signaling, and that both CLICs are required, non-redundantly, to facilitate the S1PR1-Gαi branch of the pathway and to activate Rac1.

Because the EXC-4 N-terminus promotes membrane accumulation but does not by itself rescue *C. elegans exc-4* null mutants (Berry et al., 2003; Berry and Hobert, 2006), we hypothesized that the CLIC1 and CLIC4 N-termini also primarily mediate localization, while CLIC-specific signaling functions are encoded by their C-termini. To test this, we created HA-tagged chimeras (see Methods, and Figs. 1C, 2A), where the N-terminus of CLIC1 was fused to the C-terminus of CLIC4 (C1-C4), and vice-versa (C4-C1), and assessed their expression in HUVEC, finding that the chimeras are well-expressed in these cells (Fig. 2A). We next tested the ability of these chimeras to rescue *CLIC1*^*KD*^ or *CLIC4*^*KD*^ phenotypes in HUVEC, and found that full-length CLIC1 and C4-C1, but not the C1-C4 chimera, rescued *CLIC1*^*KD*^ defects in S1P-induced Rac1 activation and endothelial barrier enhancement measured via trans-endothelial electrical resistance, or TEER (Fig. 2B, D). Conversely, *CLIC4*^*KD*^ defects were significantly rescued by full-length CLIC4 and by C1-C4, but not by the C4-C1 chimera (Fig. 2C, E). These results demonstrate that the CLIC1 and CLIC4 C-termini provide specificity to CLIC function in endothelial S1P-induced Rac1 activation.

**Fig. 2:**
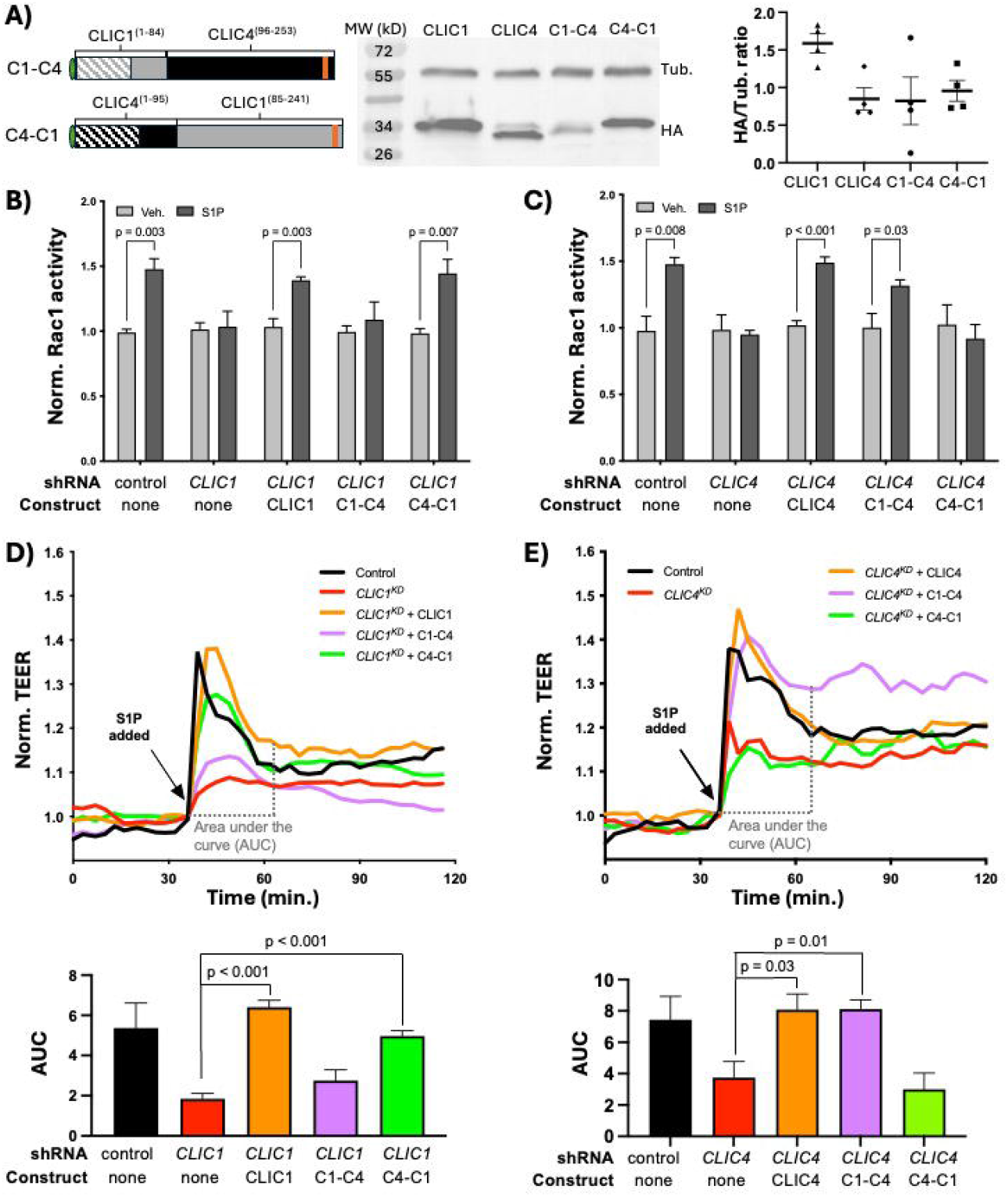
CLIC1 and CLIC4-specific functions in S1P signaling are encoded by their C-termini. (**A**) cartoon representation of the C1-C4 and C4-C1 chimeras, and western blot analysis and quantification (n=4) of their expression in HUVEC (see Methods). (**B**) full-length CLIC1 and the C4-C1 chimera rescue the Rac1 activation defect caused by *CLIC1*^*KD*^, as assessed by G-LISA (see Methods). (**C**) full-length CLIC4 and the C1-C4 chimera rescue the Rac1 activation defect caused by *CLIC4*^*KD*^, as assessed by G-LISA. In panels (B) and (C) each bar represents the mean of n≥3 replicates, the error bars represent standard deviation (hereafter, SD), and significance between vehicle control (Veh.) and S1P-treated cells was calculated via unpaired t-tests. HUVEC monolayers form a barrier whose strength can be measured via a trans-endothelial electrical resistance (TEER) assay, and stimulation by S1P rapidly, and reversibly, enhances HUVEC barrier function (Garcia et al., 2001). We previously showed that *CLIC1* or *CLIC4* knockdown reduced S1P-induced barrier strengthening (Mao et al., 2021), and here we show that (**D**) full-length CLIC1 and the C4-C1 chimera rescue the barrier defect caused by *CLIC1*^*KD*^ (see Methods), while (**E**) full-length CLIC4 and the C1-C4 chimera rescue the TEER defect caused by *CLIC4*^*KD*^. Each trace represents the mean TEER measurement of n≥3 samples, normalized to the TEER value at time of S1P addition (see Methods). To quantify these results, the area under the curve (AUC) ± standard error of the mean (SEM) from the time of S1P addition (∼30min. after beginning of resistance measurements) until time of maximal recovery to baseline in control (∼60 min. after beginning of experiment, ∼30 min. after S1P addition) was calculated for each condition (see Methods), and these results are shown in the accompanying bar graphs. Significance was calculated using unpaired 2-tailed t tests.

### The PTMD is necessary for S1P-induced CLIC4 membrane re-localization and Rac1 activation

To further probe the function of the CLIC N- and C-termini, we generated a truncated CLIC4 lacking the PTMD (hereafter (ΔPTMD)CLIC4. See Figs. 1B, 3A.) and expressed this truncated protein in HUVEC. We assessed expression via western blotting, and although we detected a band of the expected size, expression levels were consistently low for this truncation (Fig. 3A). We found that (ΔPTMD)CLIC4, like full-length protein, localized to the cytoplasm and nucleus in the absence of S1P (Fig. 3B). However, and unlike full-length CLIC4, this localization was not affected by S1P treatment (Fig. 3B). Therefore, the CLIC4 PTMD is necessary for re-localization from the cytoplasm to the plasma membrane in response to S1P. We next assessed (ΔPTMD)CLIC4 function and found that this protein did not rescue the Rac1 activation (Fig. 3C) or TEER defects (Fig. 3D) caused by *CLIC4*^*KD*^. This lack of function suggests that either the lower levels of (ΔPTMD)CLIC4 expression precludes efficient rescue, or that PTMD-mediated re-localization of CLIC4 to the plasma membrane is required for its function in S1P-induced Rac1 activation.

**Fig. 3:**
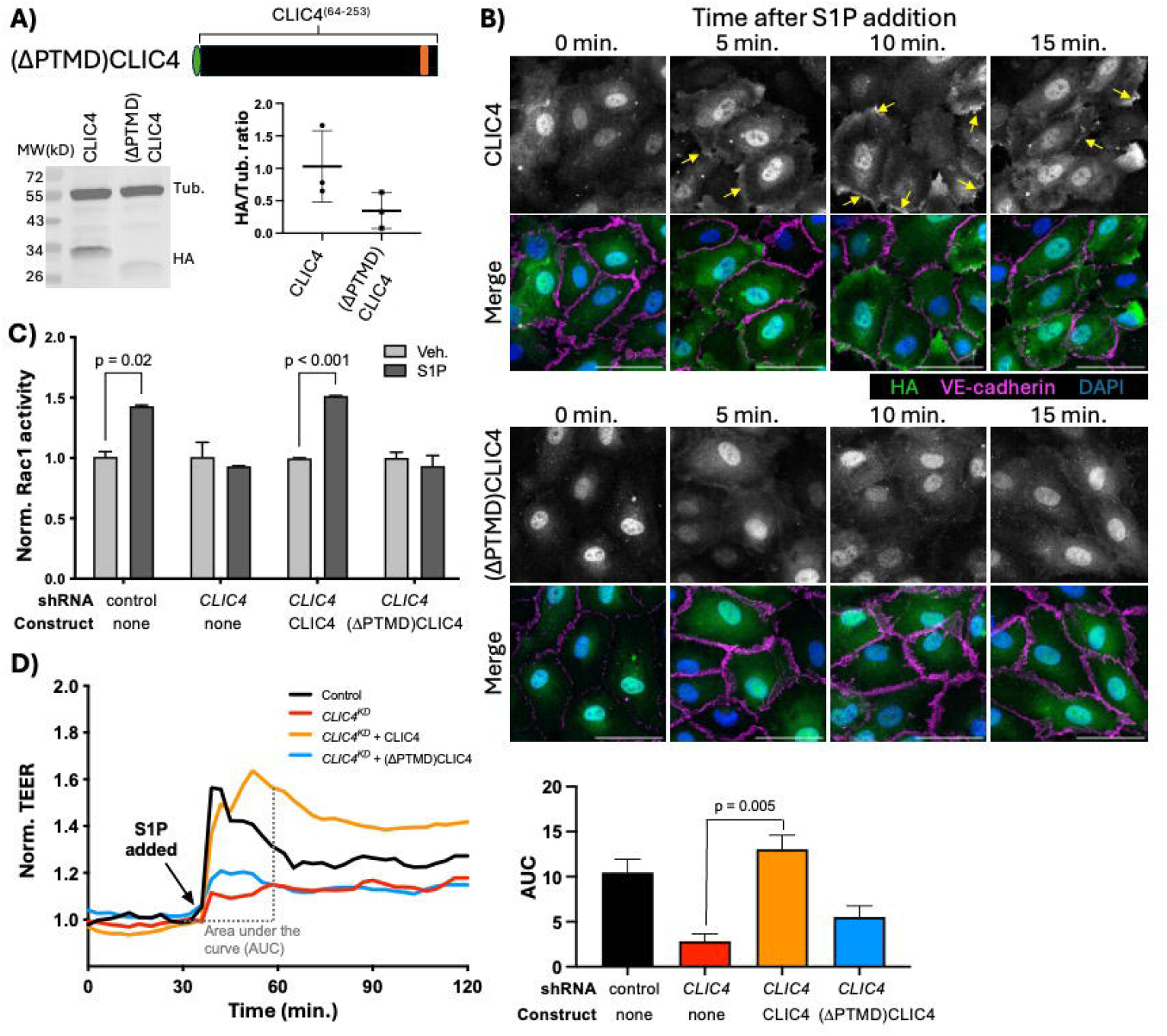
The N-terminal putative transmembrane domain (PTMD) is required for CLIC4 relocalization and Rac1 activation in response to S1P. (**A**) cartoon representation of the HA-tagged (ΔPTMD)CLIC4 construct, and western blot analysis and quantification (n=3) of its expression in HUVEC (see Methods), as compared to full-length HA-CLIC4. (**B**) Immunofluorescence of full-length HA-tagged CLIC4 (top) and (ΔPTMD)CLIC4, and their localization after treatment with S1P (see Methods). Both constructs showed nuclear and cytoplasmic accumulation prior to S1P treatment. Noticeable accumulation of full-length CLIC4 was observed near the cell cortex (outlined by VE-cadherin) within 5 minutes and remained at 10 and 15 minutes after S1P addition (yellow arrows, top rows). In contrast we did not observe similar re-localization of the (ΔPTMD)CLIC4 construct (bottom rows). Scale bar represents 50µm in all panels. (**C**) the (ΔPTMD)CLIC4 construct does not rescues the Rac1 activation defect caused by *CLIC4*^*KD*^, as assessed by G-LISA. Each bar represents the mean of n≥3 replicates, the error bars represent SD, and significance between vehicle control (Veh.) and S1P-treated cells was calculated via unpaired t-tests. (**D**) the (ΔPTMD)CLIC4 construct does not rescue the TEER defect caused by *CLIC4*^*KD*^. Each trace represents the mean TEER measurement of n≥3 samples, normalized to the TEER value at time of S1P addition, and quantitative AUC analysis of TEER calculation was carried out as described in Fig. 2D and 2E.

### A membrane-localized form of the CLIC4 C-terminus functions in S1P-induced Rac1 activation

We sought to determine if the prime function of the CLIC4 N-terminus is to facilitate plasma membrane localization, and, based upon the finding that the C-terminus of CLICs confer signaling specificity (Fig. 2), we also posited that the C-terminus may be the domain responsible for CLIC4-mediated Rac1 activation. To address these questions, we used a heterologous membrane-localization signal, the myristylation (myr) signal of the kinase Lck (Bijlmakers et al., 1997; Zlatkine et al., 1997), to induce constitutive membrane localization of ectopically expressed CLIC4. We generated two new constructs with myr signal appended to the N-terminal HA-tag: a full length myr-CLIC4, to assess effects of the myr signal on full-length CLIC4 expression and function, and a truncated myr-(ΔPTMD)CLIC4, and assessed expression of these constructs in HUVEC via western blotting, finding single bands of the expected size, with the truncated form again consistently showing lower expression than the full-length protein (Fig. 4A). Notably, both myr-CLIC4 and myr-(ΔPTMD)CLIC4 displayed constitutive membrane accumulation in the absence of S1P (Fig. 4B), indicating that, as expected, the myr signal confers constitutive membrane accumulation of both CLIC4 constructs. Importantly, both myr-CLIC4 and myr-(ΔPTMD)CLIC4 rescued the Rac1 activation (Fig. 4C) and TEER defects (Fig. 4D) caused by *CLIC4*^*KD*^. Thus, we conclude that the myr signal does not interfere with CLIC4 function. The finding that myr-(ΔPTMD)CLIC4 rescues *CLIC4*^*KD*^, leads us to conclude that the C-terminal domain carries the effector function of CLIC4 downstream of S1PR. Finally, as the N-terminus, contains sequences necessary for GST-like and ion channel activities, our findings establish that GST or ion channel activity is not necessary for CLIC4 function in S1P-induced Rac1 activation.

**Fig. 4:**
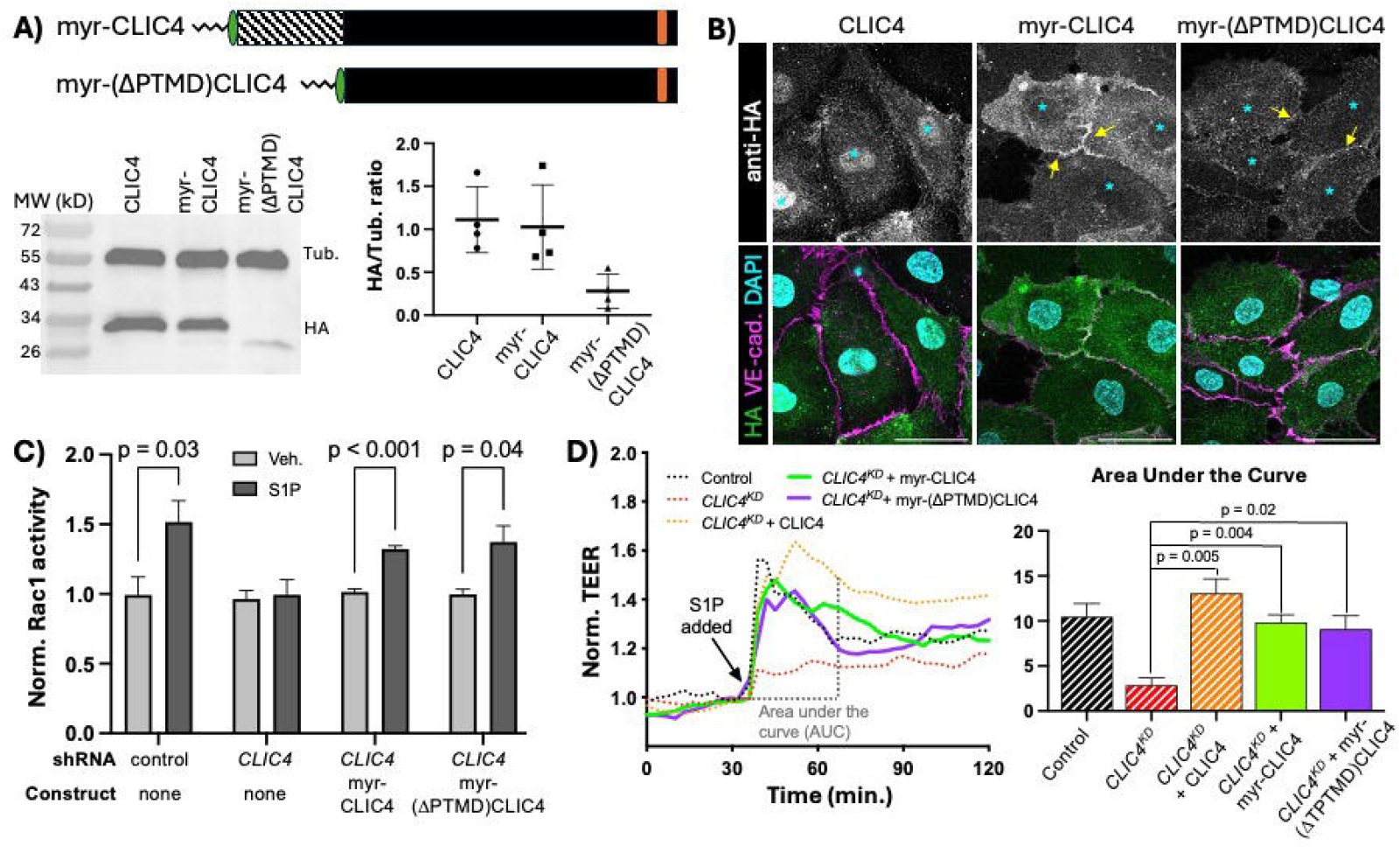
Membrane localized CLIC4 C-terminus is sufficient to promote Rac1 activation in response to S1P. (**A**) cartoon representation of HA-tagged full-length myristylated (myr) CLIC4 and myr-(ΔPTMD)CLIC4, lacking the N-terminal putative transmembrane domain. Western blot analysis and quantification (n=3) of construct expression in HUVEC (see Methods), shows lower expression of myr-(ΔPTMD)CLIC4 as compared to full-length myr-CLIC4. (**B**) Immunofluorescence showing accumulation of HA-tagged full-length CLIC4 (first column), myr-CLIC4 (second column) and myr-(ΔPTMD)CLIC4 (third column) at steady state (no S1P). Note that both myristylated constructs display accumulation at the membrane (yellow arrows), and decreased nuclear accumulation (cyan asterisks) when compared to wildtype full-length CLIC4. Scale bar represents 50µm in all panels. (**C**) both the full-length myr-CLIC4 and the myr-(ΔPTMD)CLIC4 constructs rescue the Rac1 activation defect caused by *CLIC4*^*KD*^, as assessed by G-LISA. Each bar represents the mean of n≥3 replicates, the error bars represent SD, and significance between vehicle control (Veh.) and S1P-treated cells was calculated via unpaired t-tests. (**D**) (**C**) both the full-length myr-CLIC4 and the myr-(ΔPTMD)CLIC4 constructs rescue the TEER defect caused by *CLIC4*^*KD*^. Each trace represents the mean TEER measurement of n≥3 samples, normalized to the TEER value at time of S1P addition, and quantitative AUC analysis of TEER calculation was carried out as described in Fig. 2D and 2E. Dashed lines and bars (Control, *CLIC4*^*KD*^, and *CLIC4*^*KD*^ + CLIC4) are the same data shown in Fig. 3D

## Discussion

CLICs have emerged as conserved players in Rho-family GTPase signaling in contexts as diverse as the *C. elegans* excretory canal (Arena et al., 2022), mouse kidney podocytes (Tavasoli et al., 2016), human endothelial cells and mouse vasculature (Mao et al., 2021; Kleinjan et al., 2023). The molecular function of CLICs has long-remained mysterious, with structural and biochemical studies suggesting they may function as channel and/or thiol-transferases (reviewed in Argenzio and Moolenaar, 2016). However, questions remained as to the relevance of these postulated CLIC activities for their emerging role in Rho-family GTPase signaling. We addressed this issue by undertaking structure-function studies in human endothelial cells and our results demonstrate that the signaling functions of CLIC1 and CLIC4 reside at their C-termini, while the N-terminus is dispensable for function in GPCR signaling and mainly plays a role in promoting plasma membrane accumulation upon GPCR activation. These results are consistent with findings in *C. elegans*, where it was shown that the PTMD of EXC-4, which lacks a critical cysteine residue required for GST-like activity, is necessary and sufficient for plasma membrane accumulation and is required for EXC-4 function (Berry et al., 2003; Berry and Hobert, 2006). Moreover, we also found that CLIC4 constructs lacking the N-terminal PTMD are expressed at lower levels than full-length protein, even when tethered to the plasma membrane, suggesting that the CLIC4 PTMD may encode determinants that promote stability and/or counteract degradation signals located at the C-terminus.

The activity of Rho-family GTPases is precisely controlled by regulating their localization to the plasma membrane, and by membrane localization of activators (guanine-exchange factors, or RhoGEFs) and inhibitors (GTPase-activating proteins, or RhoGAPs). Therefore, our finding that plasma membrane accumulation of the CLIC4 C-terminus is required for Rac1 activation in response to S1P suggests a model where CLIC C-termini activate Rho-family GTPases either directly or as part of a complex with RhoGEFs and/or RhoGAPs. If CLICs directly activate Rho-family GTPases, we might expect that constitutive recruitment of full-length, or just the C-terminus of, CLIC4 to the plasma membrane could be sufficient to activate Rac1. However, we did not observe activation of Rac1 in cells expressing myr -CLIC4 or myr-(ΔPTMD)CLIC4 in the absence of S1P (Supplemental Fig. S1), indicating that CLICs are necessary, but not sufficient, to activate Rho-family GTPases in response to GPCR ligands. We expect that continued structure-function studies to elucidate C-terminal sequences required for EXC-4/CLIC function, combined with discovery of EXC-4/CLIC-interacting proteins required for function, will reveal the novel and conserved molecular mechanisms by which CLICs regulate Rho-family GTPase activity.

## Materials and Methods

### Primary cells and cell culture

Pooled HUVECs from different donors were either directly isolated from human umbilical cords following established protocols (Jaffe et al., 1973) or purchased (Lonza, Cat# C2519A). HUVEC were grown in EGM-2 Endothelial Cell Growth Media (Lonza), including all supplements provided, on culture dishes coated with rat tail type I collagen (Corning). 293T cells were acquired from the American Type Culture Collection and maintained in high-glucose Dulbecco’s modified Eagle’s medium (Gibco) with 10% heat-inactivated fetal bovine serum (HI-FBS) and 0.01% penicillin-streptomycin. Unless otherwise noted, cells were cultured under standard conditions in a humidified incubator at 37°C, 5% CO_2_.

### RNAi-mediated CLIC1 and CLIC4 knockdown constructs

Validated control (scrambled) shRNA, and human CLIC1 and CLIC4 shRNA-expressing lentiviruses, based on the pLKO.1 vector backbone (Moffat et al., 2006), from Sigma-Millipore were as previously-described (Tung et al., 2009; Tung and Kitajewski, 2010; Mao et al., 2021; Kleinjan et al., 2023).

### Tagged CLIC1 and CLIC4 constructs

Full-length HA-tagged CLIC1 and CLIC4 plasmids were previously described (Mao et al., 2021), and the derived chimeric and truncated variants used here were made using standard molecular cloning techniques, and DNA sequencing was performed to verify the correct sequence before experiments (details available upon request). All HA-tagged constructs were cloned into the pCCL lentivirus vector (Dull et al., 1998) for delivery and expression in HUVEC

### Lentivirus-mediated stable overexpression and/or knockdown in HUVEC

To perform knockdown and/or overexpression in HUVEC, a lentiviral infection system was utilized. For lentiviral gene transfer, 293T cells were transfected using the calcium phosphate approach with the following combination of plasmids: 3μg of pVSVG, 5μg of pMDLg/pRRE, 2.5μg of pRSV-Rev, and 10μg of the overexpression (pCCL-based) and/or knockdown (pLKO.1-based) vectors of interest. Transfected 293T cells were allowed to produce lentivirus and the supernatant was collected 48 hours post transfection. The supernatant collected was passed through a 0.45μm filter and then added onto HUVEC. A single round of infection was performed for 24hrs. The primary cells were allowed to express shRNA or overexpression constructs for at least 48 hours before experiments. A red-flourescent construct (pCCL-RFP) was included in all experiments to assess infection efficiency.

### Reagents and Antibodies

Sphignosine-1-phosphate (S1P) was obtained from Enzo Life Sciences (BML-SL140). Antibodies to human CLIC1 (Abcam ab28722, Rabbit, 1:250), human CLIC4 (Novus Biologicals NBP1-85574, Rabbit, 1:250), HA (GenScript A01244, Mouse, 1:500), tubulin (Sigma-Aldrich T6074, 1:5000), and VE-cadherin (Abcam ab33168, Rabbit, 1:250) were used for immunoblotting and immunofluorescence as described below.

### Immunoblotting

HUVECs were washed with cold PBS, and lysates were collected with TENT lysis buffer (50mM Tris pH 8.0, 2mM EDTA, 150mM NaCl, and 1% Triton X-100) containing a protease inhibitor cocktail (EMD Chemicals Inc.). Protein lysates were collected by centrifugation at 14000rpm for 10 minutes. Lysates were boiled at 95°C for 5 minutes with addition of sample buffer containing SDS and β-mercaptoethanol. Protein concentrations were measured using the Bradford protein assay (Bio-Rad Laboratories). Volumes were adjusted to ensure equal amounts of protein loading. SDS-PAGE was performed for 1 hours at 150V, followed by wet transfer of proteins onto a nitrocellulose membrane for 1 hour at 100V. Blocking of the nitrocellulose membrane was with 5% BSA in TBS-Tween solution for 1 hour. Primary antibody incubation was done in 2.5% BSA overnight at 4°C, and secondary antibody incubation was done the next day for 2hr at room temperature. The membrane was developed using Clarity™ Western ECL Substrate (Bio-Rad) and protein bands were observed. Densitometry analysis was performed with GelAnalyzer 19.1 by Istvan Lazar Jr., PhD and Istvan Lazar Sr., PhD, CSc (available at www.gelanalyzer.com).

### Immunofluorescence

Cells were plated on 8-well collagen-coated chamber slides (Ibidi). 50,000 cells were seeded onto each well overnight. The next day cells were serum starved for 3 hours prior to S1P treatment. 1μM S1P for various times points was used to treat HUVECs, followed by fixation with 4% PFA for 10 minutes. Cells were then washed 3 times with PBS and blocked with 3% BSA and 0.1% TritonX-100. Primary antibodies were used at the listed dilutions in blocking solution and incubated overnight at 4°C. Cells were then washed with 1XPBS, 3 times for 5 minutes each and incubated with secondary antibody Alexa Fluor 488 (green) or Alexa Fluor 647 (far-red) at 1:1000 in blocking solution for 2hrs. After washing cells 3 times after secondary antibody incubation, cells were mounted using VectaShield with DAPI (Vector Labs). Slides were imaged using Airyscan confocal microscopy with a Zeiss laser scanning microscope (LSM800). Images analysis was done with ZEN software under the same acquisition setting for among all cell lines in each experiment.

### Rac1 G-LISA activation assay

Assays were performed using a Rac1 (Cytoskeleton, BK128) G-LISA Activation Assay Kits. HUVECs were serum-starved in EGM-2 with 1% serum overnight and with serum-free EBM-2 for an additional 3 hours the following day. The cells were then stimulated with 1 μM S1P for Rac1 activation. Cell lysates were harvested and snap-frozen in liquid nitrogen. The assay was then performed based on the manufacturer’s protocol.

### Trans-endothelial electrical resistance (TEER) assay

An ECIS array plate (Applied Biophysics) containing circular 250μm diameter active electrodes connected in parallel on a common gold pad was coated with rat tail type I collagen (Corning). HUVEC cells were seeded at 50,000 cells per well and allowed to grow overnight. Cells were serum starved (EBM-2, Lonza) for 2 hours, followed by a 30-min baseline resistance stabilization with the Electrical Cell-Substrate Impedance Sensing (ECIS) system (Applied Biophysics, model 1600R). 1μM S1P or BSA vehicle control were administered, and trans-endothelial resistance was monitored at a frequency of 4000Hz with measurements taken at 3-min intervals for 24 hours. Quantifications were performed by area under the curve analysis using GraphPad Prism software.

### Statistics

All experiments were repeated at least three times, and statistical tests were performed using GraphPad Prism software. Area under the curve averages and standard error of the mean (SEM) were computed using the Gagnon method, which produces a single value, so individual AUC measurments for each replicate are are not determined. Unless otherwise noted, unpaired 2-tailed t tests were used, and P values less than 0.05 were considered statistically significant.

## Supporting information

Mao et al, Supp Fig. 1

## Author Contributions

Conceptualization: DYM, JJJ, DDS, JK

Data curation: DYM, JJJ

Formal analysis: DYM, JJJ, DDS

Funding acquisition: JJJ, DDS, JK

Investigation: DYM, JJJ

Methodology: DYM, JJJ, DDS, JK

Project administration: DDS, JK

Resources: DDS, JK

Supervision: DDS, JK

Validation: DYM, JJJ

Visualization: DYM, JJJ, DS

Writing – original draft: DYM, JK

Writing – review & editing: DYM, JJJ, DS, JK

## Acknowledgments

The authors would like to thank Alison Kitajewski for technical support, Matthew J. Kleinjan for help with early stages of this work, and members of the Kitajewski and Shaye laboratories for fruitful discussions during this project.

## Supplementary Material

### Data Availability Statement

The datasets and reagents generated for this study will be made available by the authors upon request, without undue reservation.

