## Supplementary figures and images for "Chloride intracellular channel (CLIC) protein function in S1P-induced Rac1 activation requires membrane localization of the C-terminus, but not thiol-transferase nor ion channel activities"

### Mao et al, Supp Fig. 1

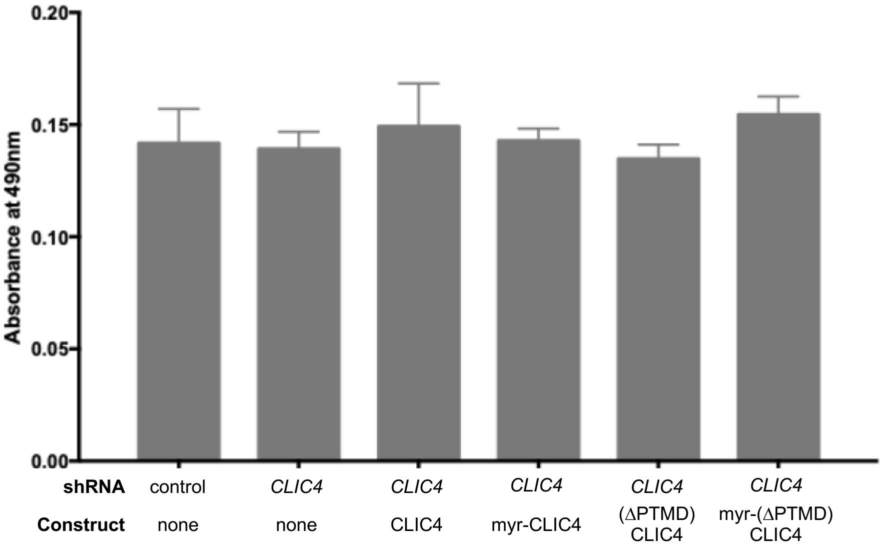


**Supplemental Figure S1:** Basal Rac1 activity is not increased by membrane tethered CLIC4
